# Characterization of the microbiome of *Aedes albopictus* populations in different habitats from Spain and São Tomé: implications for vector control

**DOI:** 10.1101/2024.02.24.581876

**Authors:** Tiago Melo, Carla Alexandra Sousa, Sarah Delacour-Estrella, Daniel Bravo-Barriga, Gonçalo Seixas

## Abstract

The mosquito microbiome significantly influences vector competence, including in *Aedes albopictus*, a globally invasive ectoparasite. Describing the microbiome and *Wolbachia* strains of *Ae. albopictus* from different regions can guide area-specific control strategies. Mosquito samples from Spain and São Tomé were analyzed using 16S rRNA gene sequencing and metagenomic sequencing. *Wolbachia* infection was genotyped, and statistical analysis compared infection patterns by sex and population. Results revealed a dominance of dual *Wolbachia* infections in the microbiome of both populations of *Ae. albopictus*, especially among females. The two populations presented a shared microbiome, although 5 and 9 other genera were only present in Spain and São Tomé populations, respectively. Genera like *Pelomonas* and *Nevskia* were identified for the first time in *Aedes* mosquitoes. This study is the first to describe the *Ae. albopictus* bacteriome in Spain and São Tomé, offering insights for the development of targeted mosquito control strategies.

## Introduction

Vector-borne diseases, which account for 20% of infectious diseases worldwide, pose a significant global health threat. Illnesses like dengue fever, yellow fever, chikungunya, and Zika affect nearly 3.9 billion people (WHO, 2022b) and effective management and control of these disease vectors are crucial for reducing these risks of transmission.

*Aedes albopictus*, a mosquito species known for its ability to transmit diseases and nuisance to humans (Capinera, 2008), has a worldwide distribution (Manni et al., 2017) and is responsible for spreading dengue, chikungunya, and Zika viruses (Effler et al., 2005; Paupy et al., 2009; Ramchurn et al., 2009). Its invasive nature (Medlock et al., 2015) and adaptability to different environments (Swan et al., 2022) make it a significant threat to human health. Originally from Southeast Asia, *Ae. albopictus* has expanded its range to other continents in a very short period, facilitated by global trade of used tires, ornamental plants, and road traffic (Gratz, 2004; Paupy et al., 2009; Eritja et al., 2017). Moreover, in recent decades, rapid urbanization has contributed to proliferation of *Ae. albopictus* populations, and this, added to inadequate or absent vector control measures, have further increased the risk of pathogen transmission by this species (Hasan et al., 2016). The unstoppable dispersion of *Ae. albopictus* mosquitoes in São (Reis et al., 2017) Tomé and Principe and Spain (Goiri et al., 2020), highlights the need of understanding and managing this species in these regions. São Tomé and Principe, an island nation in the Gulf of Guinea, is particularly vulnerable to vector-borne diseases due to its tropical climate and limited healthcare infrastructure (WHO 2022b) but colonization events should be rare but colonization events should be rare (Loiseau et al., 2019). *Aedes albopictus* was first identified in São Tomé in 2016, although it is probable that the species had inhabited the island for a decade prior to this discovery (Reis et al., 2017). In Spain, the first *Ae. albopictus* was detected in 2004, at Sant Cugat del Vallès (Aranda et al., 2006). Since then, the mosquito’s presence has proliferated across the country (Goiri et al., 2020) due to multiple introductions from abroad (Lucati et al., 2022). This expansion has escalated concerns among public health officials, emphasizing the need for increased surveillance and control measures (ECDC, 2019; Monge et al., 2020; Navero-Castillejos et al., 2021).

The host physiology of *Aedes* mosquitoes is heavily influenced by their microbiota, which has been shown to impact various aspects of reproduction, egg production, blood digestion (Gaio et al., 2011; Coon et al., 2016; Sicard et al., 2019), regulation of immunity (Xi et al., 2008), genetic diversity, vector competence, and host-pathogen interactions (Dickson et al., 2017; Souza-Neto et al., 2019; Lucati et al., 2022). So, innovative vector control strategies have increasingly focused on manipulating the mosquito microbiome to reduce their capacity for disease transmission (Jupatanakul et al., 2014) and control of populations. One promising approach involves introducing exogenous *Wolbachia* strains into *Ae. albopictus* populations, which has shown potential in disrupting pathogen transmission (Blagrove et al., 2013). However, most of the *Ae. albopictus* populations are already naturally infected with this endosymbiont (Ahmad et al., 2017) and different *Wolbachia* strains may exhibit various interactions when coexisting within a host (Blagrove et al., 2013). Describing the native strains of our target-populations is the first step to assess possible interactions between the naturally occurring *Wolbachia* strains and the introduced type.

The genetic background of any given species plays a role in shaping its microbiome (Novakova et al., 2017). Different populations of mosquitoes may have genetic variations that affect their immune responses, susceptibility to certain microorganisms, and overall interactions with the microbiome. In the same way, different geographical locations may have unique microbial communities in soil, water, and vegetation, and the local climate and temperature can impact both the mosquito and the microorganisms it harbors (Dada et al., 2021). Given the significance of understanding and managing vector-borne diseases, this study aimed to examine and to compare the microbiome composition and characterize the presence of *Wolbachia* strains in populations of *Ae. albopictus* from two countries with distinct climatic, habitat, epidemiological, and geographical conditions: São Tomé and Príncipe (Africa) and Spain (Europe). Investigation on the composition of the microbiome and *Wolbachia* strains in *Ae. albopictus* populations from these distinct geographical regions could provide valuable knowledge into potential control measures tailored to the specific problems of each area. This information could help develop new vector control strategies and contribute to reduce the global burden of vector-borne diseases.

## Materials and Methods

### Mosquito collection and rearing

Eggs were collected using ovitraps installed in three distant regions of Spain (Paterna, Monzón, and Rincón de La Victoria) and one region of São Tomé (Figure 1) between late 2021 and early 2022, with the assistance of collaborators. Four *Ae. albopictus* colonies were subsequently established and maintained at the high-security insectary of the “In Vivo Arthropod Security Facility” (VIASEF) available at IHMT. Upon hatching in dechlorinated water, larvae were reared under controlled laboratory conditions (temperature: 26 ± 2°C, relative humidity: 70 ± 5%, photoperiod: 12h/12h light/dark) and fed with fish food. Adult mosquitoes were provided with a 10% glucose solution, and females were blood-fed on *Mus musculus* two to three times a week. The process of handling the animals used occurred under supervision and was carried out based on Community Council standards European Union of 24 November 1986 (86/609/EEC) and national legislation in force (Decree-Law 129/92 of June 2nd, Ordinance No. 100/92 of October 23rd).

**Figure 1.**
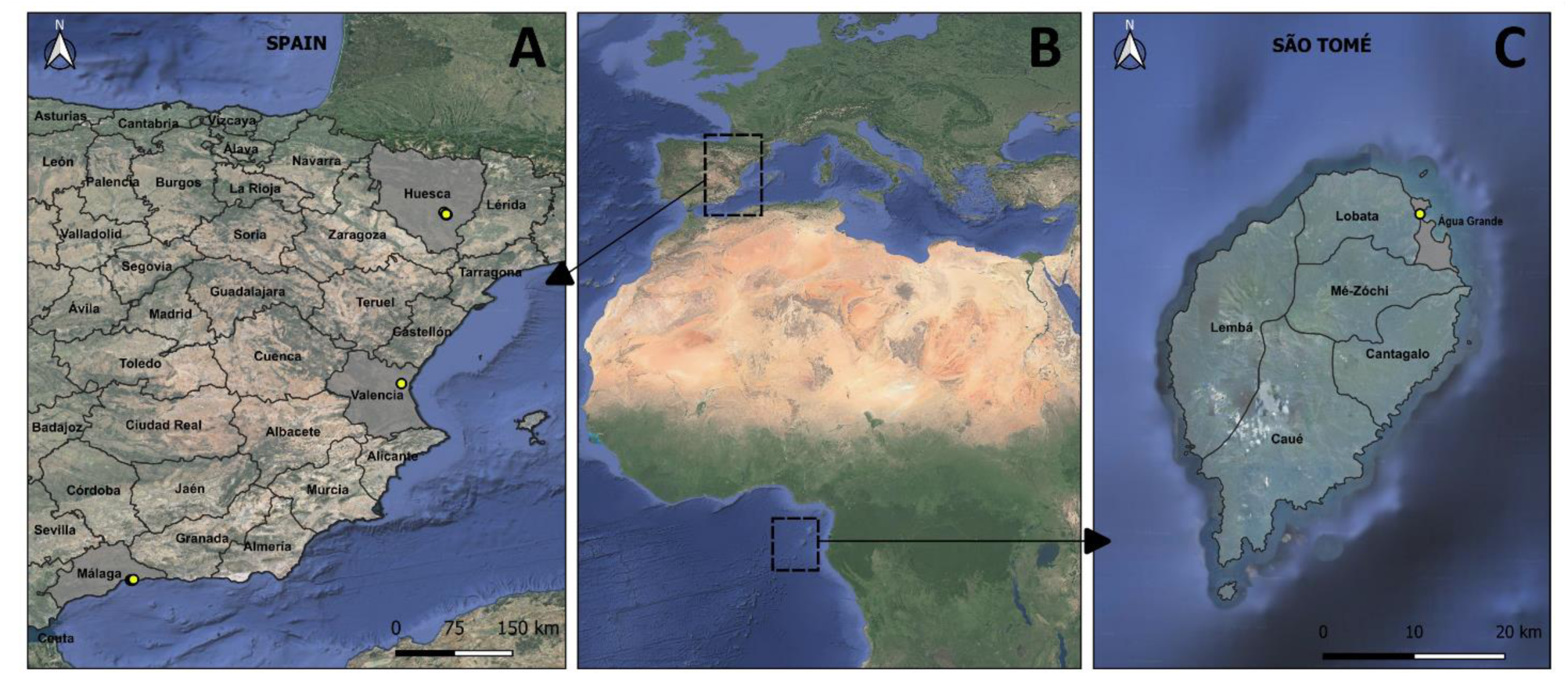
– Map indicating collection sites for *Ae. albopictus*, per continent (B): (A) Spain and (C) São Tomé.

### DNA extraction

DNA extractions (N=199) were carried out using different methods based on the sample type. For 16S rRNA gene sequencing (N=19), the “NZY tissue gDNA isolation kit” (NZYtech, Portugal) was used. For the remaining samples (N=180), ribosomal DNA extraction protocol adapted from (Collins et al., 1988) was employed. Negative controls were included during DNA extraction to rule out any potential contamination.

### 16S Metagenomic sequencing

The bacterial composition of *Ae. albopictus* microbiota was investigated by sequencing the V3 and V4 regions of the bacterial 16S rRNA gene in 19 samples. Library construction, sequencing, and bioinformatics analysis were performed by Eurofins Genomics Europe Sequencing GmbH (Konstanz, Germany). All sequences have been submitted to NCBI under the accession number for SRA data PRJNA1028981 (temporary submission ID: SUB13907152). SRA records will be accessible with the following link after the release date (2024-11-01): http://www.ncbi.nlm.nih.gov/bioproject/1028981.

### *Wolbachia* genotyping

PCR genotyping of *Wolbachia* infection in 180 samples used *Wolbachia*-specific primers 328Fw and 691Rv for *w*AlbA, and 183F and 691R for *w*AlbB. PCR mix and thermal cycling conditions were standardized. Universal primers *w*sp 81Fw and 691Rv were used to confirm *Wolbachia*-negative samples. Primers and PCR conditions details are provided in Supp. Table S1.

### Statistical analysis

Mixed and single infection rates were calculated by sex and population (Vassarstats, n.d.). Comparisons between mixed and single infections in females and males were performed using the chi-square test. A *p*-value less than 0.05 was considered statistically significant.

## Results

### Microbiome composition

To analyze the bacterial communities within *Ae. albopictus* from Spain and São Tomé, we conducted 16S rRNA gene sequencing of the V3-V4 hypervariable regions across 19 samples. After pre-processing and quality control, five samples from São Tomé yielded 606,801 clean sequences, and 14 samples from Spain yielded 798,002 clean sequences (Table 1). All these clean sequences were classified into OTUs (Operational Taxonomic Units): 344 from São Tomé and 517 from Spain (Table 1). The OTUs represented different taxonomic levels, and a correction was made to the species found (Angly et al., 2014) (Supp Tables S2 e S3). The Figure 2 shows shared and non-shared bacterial genera among the populations.

**Figure 2.**
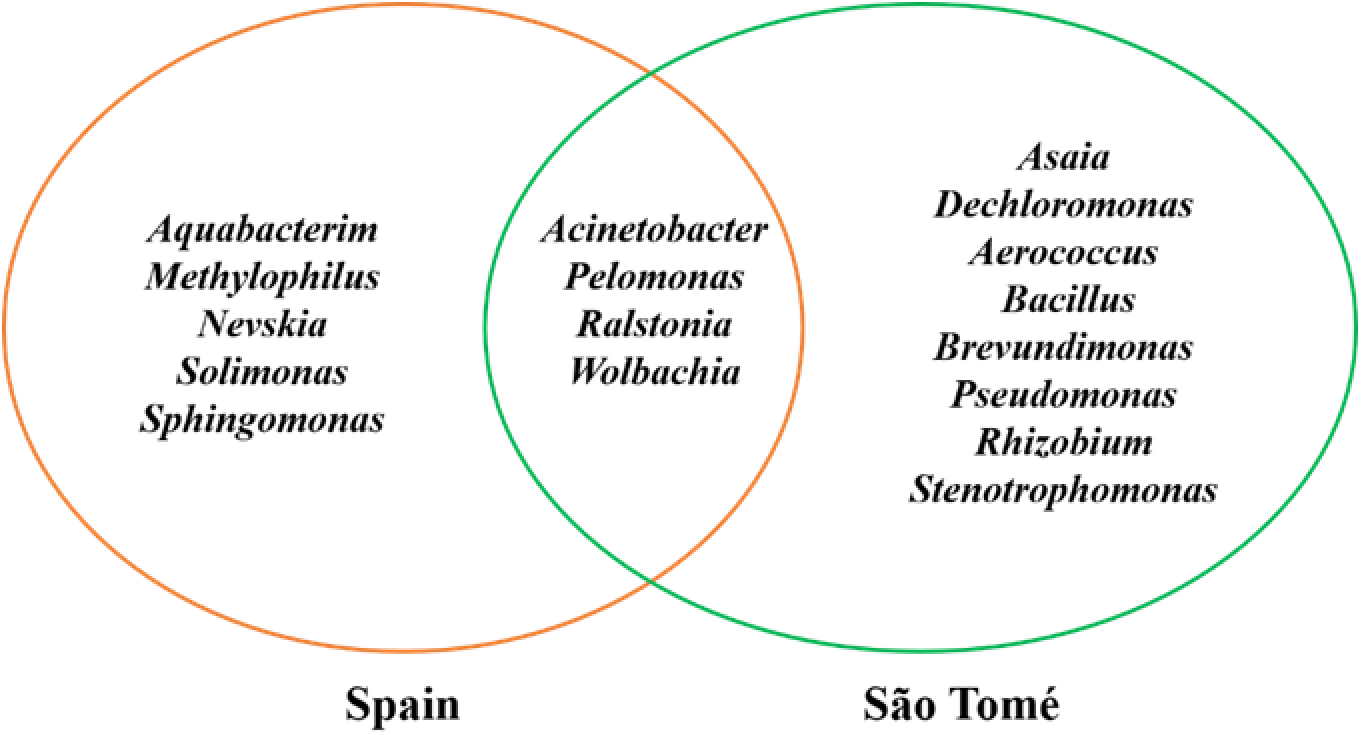
– Venn diagram showing shared and unshared genera between the Spain and São Tomé populations.

**Table 1.**
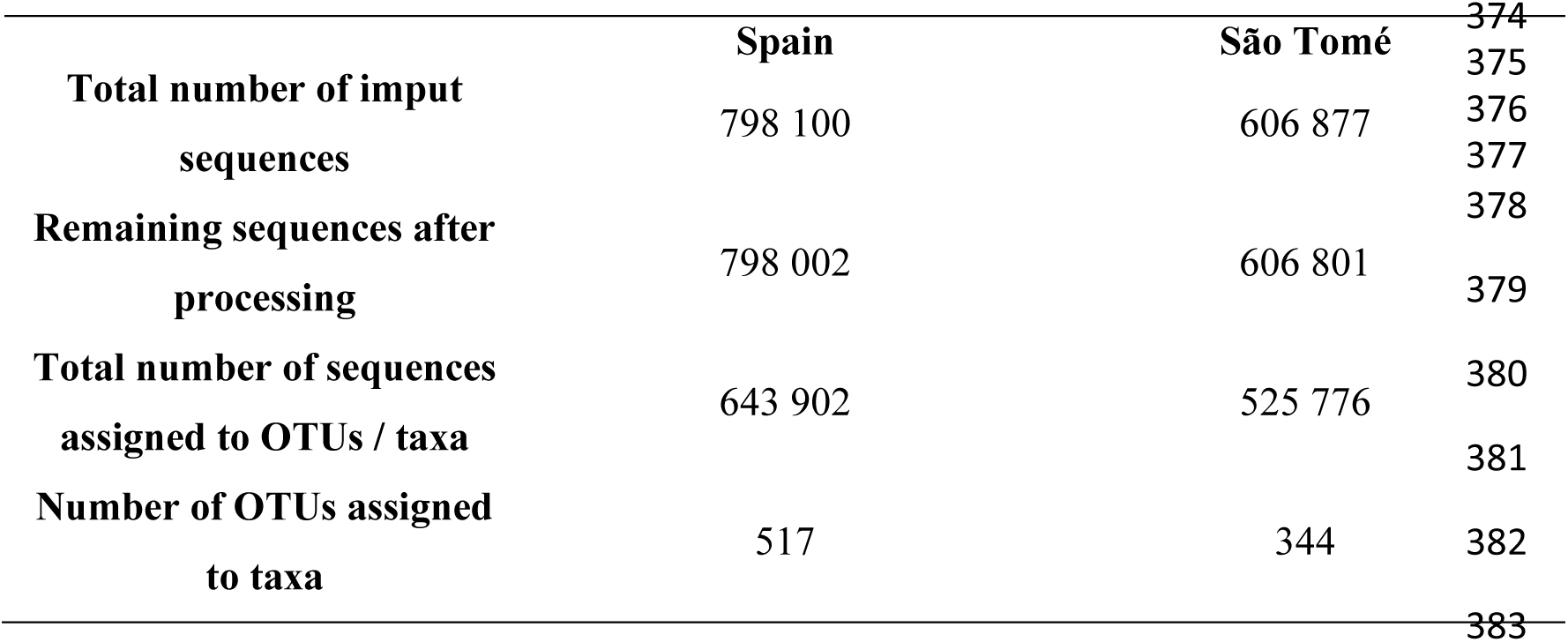
– Summarized statistics of Spain and São Tomé samples.

|  | **Spain** | **São Tomé** |
| --- | --- | --- |
| **Total number of input sequences** | 798 100 | 606 877 |
| **Remaining sequences after processing** | 798 002 | 606 801 |
| **Total number of sequences assigned to OTUs / taxa** | 643 902 | 525 776 |
| **Number of OTUs assigned to taxa** | 517 | 344 |

### Taxonomic units overview and microbiome diversity analysis

Tables 2 e 3 provide an overview of the taxonomic units identified in samples from São Tomé and Spain. Taxonomic units with readings below 0.1% were categorized as “other.” OTU readings analysis showed a predominance of the genus *Wolbachia* in all samples, ranging between 92.4% and 98.8% in São Tomé samples and between 96.1% and 97.5% in Spanish samples.

**Table 2.** – Summary of the taxonomic composition of samples from São Tomé.

| **Sample** | **Taxonomic Unit** | **OTUs** | **Reads** | **Raw Fraction** |
| --- | --- | --- | --- | --- |
| **STPF0.1** | *Wolbachia* | 191 | 101778 | 98,1% |
|  | Gammaproteobacteria | 1 | 157 | 0,2% |
|  | Other |  |  | 1,7% |
| **STPF0.2** | *Wolbachia* | 185 | 70626 | 92,4% |
|  | *Asaia* sp. | 1 | 1311 | 1,8% |
|  | Gammaproteobacteria | 3 | 1035 | 1,4% |
|  | Enterobacteriaceae | 1 | 876 | 1,2% |
|  | Erwiniaceae | 1 | 242 | 0,3% |
|  | Enterobacterales | 2 | 195 | 0,3% |
|  | *Acinetobacter* | 2 | 166 | 0,2% |
|  | *Dechloromonas* | 1 | 95 | 0,1% |
|  | Other |  |  | 2,3% |
| **STPF0.3** | *Wolbachia* | 146 | 126113 | 98,8% |
|  | Other |  |  | 1,2% |
| **STPF0.4** | *Wolbachia* | 202 | 97713 | 98,2% |
|  | Other |  |  | 1,8% |
| **STPF0.5** | *Wolbachia* | 206 | 73196 | 96,7% |
|  | *Pelomonas* | 1 | 676 | 0,9% |
|  | *Brevundimonas* | 1 | 170 | 0,2% |
|  | *Pseudomonas* | 1 | 75 | 0,1% |
|  | Other |  |  | 2,1% |

**Table 3.**
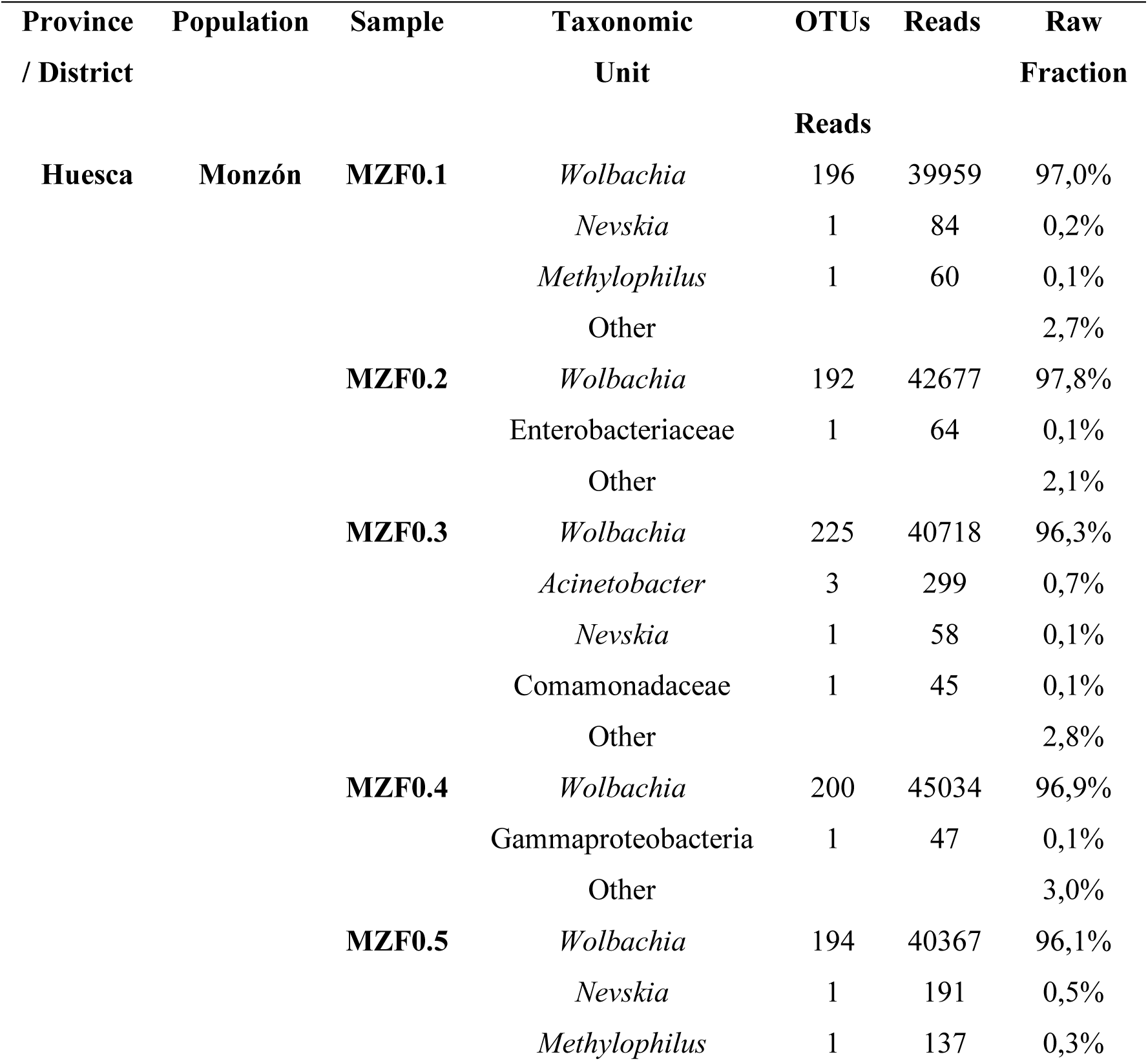
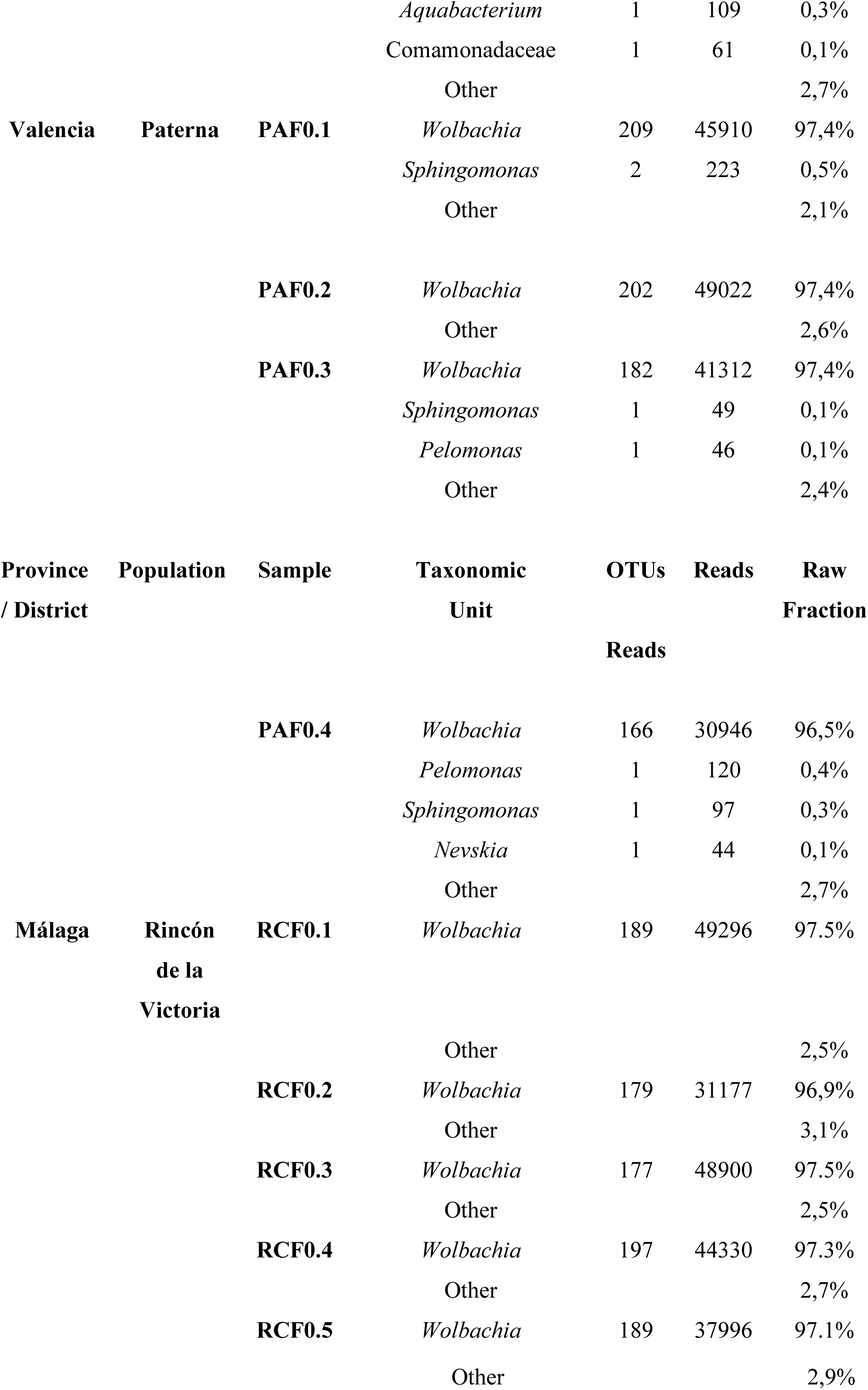
– Summary of the taxonomic composition of samples from Spain.

Shannon and Simpson indices (Table 4) revealed similar bacterial diversity between Spanish and São Tomé populations, as well as among individual samples (Supp. Table S4).

**Table 4.**
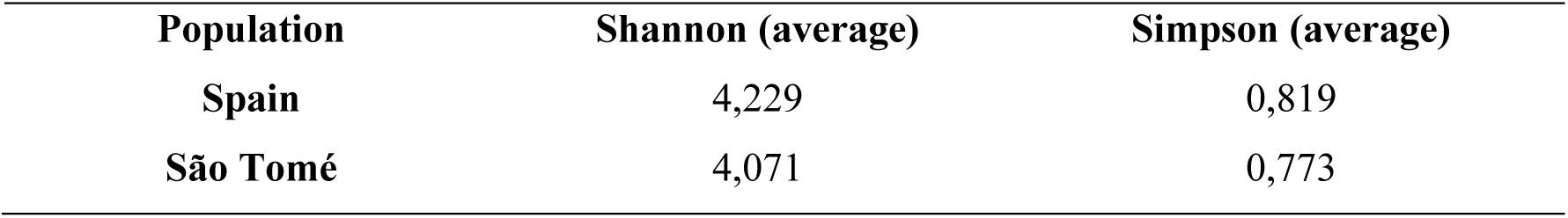
–. Average of the Shannon and Simpson indices of the Spain and São Tomé populations.

### *Wolbachia* genotyping

Of the 180 adult mosquitoes analyzed (97 females and 83 males), 178 (98.89%) were infected with *Wolbachia*. Five (2.78%) were infected with the *w*AlbA strain and 34 (18.88%) with the *w*AlbB strain. Mixed infections with *w*AlbA and *w*AlbB strains occurred in 77.22% of the infected mosquitoes (Table 5). Single *w*AlbA infections were exclusive to females (*p* < 0.05), while single *w*AlbB infections occurred in both males and females but were more prevalent in males (*p* < 0.05). Mixed infections were more common in females than males (85.57% in females and 67.47% in males, of the total number of infected mosquitoes) (*p* < 0.05). A predominance of mixed infections was observed in all locations, with rates ranging from 62.50% to 82.76%. However, simple *w*AlbB infections were detected only in the three provinces of Spain (Valencia, Monzón, and Málaga) with rates ranging from 13.79% to 31.25% (Table 6) and simple *w*AlbA infections were found only in two provinces of Spain (Valencia and Monzón). In São Tomé, only mixed infections by *w*AlbAeB were found, all in females. Furthermore, it was the only place where negative samples (N=2) were detected in males.

**Table 5.**
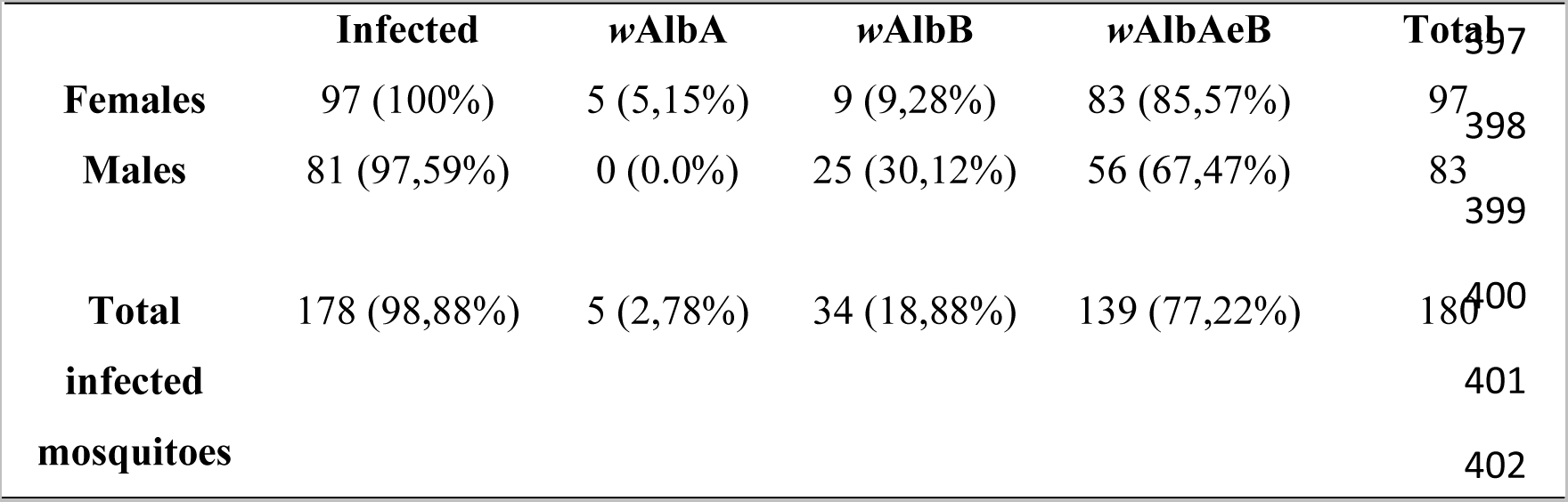
– Rate of *Wolbachia* infection in *Ae. albopictus* by sex.

**Table 6.** – Rate of *Wolbachia* infection, *w*AlbA and *w*AlbB, by locality.

| Country | Province / District | County / Region | Total | Infected | wAlbA | wAlbB | wAlbAeB |
| --- | --- | --- | --- | --- | --- | --- | --- |
| Spain | Valencia | Paterna | 16 | 16 (100%) | 1 (6,25%) | 5 (31,25%) | 10 (62,50%) |
|  | Huesca | Monzón | 116 | 116 (100%) | 4 (3,45%) | 16 (13,79%) | 96 (82,76%) |
|  | Málaga | Rincón de la Victoria | 41 | 41 (100%) | 0 (0,00%) | 13 (31,71%) | 28 (68,29%) |
| São Tomé | Água Grande | São Tomé | 7 | 5 (71,43%) | 0 (0,00%) | 0 (0,00%) | 5 (71,43%) |

## Discussion

### Aedes albopictus microbiome

*Aedes albopictus* mosquito presents a significant public health concern due to its exceptional ability to transmit emerging human pathogens and its capacity for invasive spread (Effler et al., 2005; Paupy et al., 2009; Ramchurn et al., 2009), encompassing Europe (ECDC, 2023) and Africa (Kampango & Abílio, 2016; Bennouna et al., 2017; WHO, 2022a). Microbiome play a pivotal role in shaping mosquito biology (Wang et al., 2018). So, recent attention has been drawn to the manipulation of mosquito microbiomes as a promising avenue for environmentally friendly and innovative control strategies (Scolari et al., 2019). Specifically, the bacterial component of the mosquito’s gut microbiome exerts influence on various physiological aspects, including vector competence (Jeffries et al., 2018).

This surge in interest regarding mosquito microbiome research resulted in extensive data on diverse mosquito species from various geographical locations and their respective habitats (Gratz, 2004; Medlock et al., 2015; Hasan et al., 2016; Swan et al., 2022). Nevertheless, this study represents the inaugural attempt to provide a comprehensive insight into the composition of the *Ae. albopictus* bacterial microbiome in wild populations originating from Spain and São Tomé through the sequencing of the V3-V4 regions of the 16S rRNA gene. The main phyla identified in the *Ae. albopictus* bacterial microbiome were Proteobacteria and Firmicutes, consistent with previous studies (Bennett et al., 2019; Zhao et al., 2022). Both populations exhibited unique bacterial genera (Spain: *Aquabacterium*, *Methylophilus*, *Nevskia*, *Solimonas* and *Sphingomonas*; São Tomé: *Asaia*, *Dechloromonas*, *Aerococcus*, *Bacillus*, *Brevundimonas*, *Pseudomonas*, *Rhizobium* and *Stenotrophomonas*), suggesting a geographic influence on the microbiome (Zhao et al., 2022). A shared bacterial microbiome was also observed, comprising *Acinetobacter*, *Pelomonas*, *Ralstonia*, and *Wolbachia* genus. In addition to *Wolbachia,* these results suggest the existence of a bacterial microbiome shared among populations of *Ae. albopictus* geographically very distant (Scolari et al., 2019). The presence of *Pelomonas* and *Nevskia* genera in the *Aedes* genus is a novel finding, and their role in the microbiome of hematophagous insects remains unknown, and further studies are needed to understand their interactions with hosts (Vicente et al., 2013; Zhukova et al., 2017; Santos-Garcia et al., 2020; Tabbabi et al., 2022).

Consistent with prior research, the bacterial composition of *Ae. albopictus* was predominantly governed by the *Wolbachia* genus (Lin et al., 2021). When *Wolbachia* infection is highly dominant, it can “mask” the DNA of other bacteria present (Minard et al., 2014; Bennett et al., 2019) leading to undetected and unsequenced sequences. Future research may consider individual tissue-based microbiome analysis (Rosso et al., 2018), to ensure that *Wolbachia*, which is naturally abundant in the ovaries, does not interfere with the sequencing data.

This highly dominant infection by *Wolbachia* in both populations may be related to interactions between microbiome components, such as interspecific competition for resources (Brinker et al., 2019; Scolari et al., 2019) but also to laboratory rearing conditions (Lin et al., 2021). Research into the microbiota of laboratory and wild *Aedes* mosquitoes has revealed that both are primarily characterized by a limited number of phyla (Scolari et al., 2019). Nonetheless, although the overall microbiota composition is comparable in laboratory-reared and wild mosquitoes, it was observed that the diversity of midgut bacterial communities was more extensive in mosquitoes collected from natural environments. In fact, the diversity of female midgut bacteria in *Ae. albopictus* from the laboratory was reduced when compared to females from the field (Scolari et al., 2021). These disparities could be originated from variations in the origin of the blood meal, dietary factors, and habitat (Scolari et al., 2021; Tuanudom et al., 2021). Furthermore, recent studies have indicated a more consistent and greater bacterial diversity in breeding water compared to larvae and adults, regardless of sample source, with a notable decrease in the microbial community diversity between the larvae and newly emerged adult mosquitoes (Scolari et al., 2021). The impact of blood meals and rearing within an insectary environment on the *Ae. albopictus* microbiome warrants thorough investigation and future studies should consider the analysis of larvae’s microbiome and the bacterial diversity of the breeding water.

### *Wolbachia* genotyping

To better understand the *Wolbachia* populations naturally present in these *Ae. albopictus*, A and B strains (Tortosa et al., 2010; Ahmad et al., 2017) were genotyped. Both strains were found to infect the Spanish and São Tomé populations in three different scenarios: mixed infection with both strains, simple infection with *w*AlbA, or simple infection with *w*AlbB. Mixed infection, which is predominant worldwide (Lo et al., 2002; Shaikevich et al., 2019; Hu et al., 2020; Puerta-Guardo et al., 2020; Bueno-Marí et al., 2023) was also the predominant form in this study. Several factors could explain these results, including species susceptibility to infection, facilitation of secondary infections by an active *Wolbachia* infection, or the stable maintenance of dual infections in hosts (Werren et al., 1995).

In line with previous studies (Ahmad et al., 2017; Belo, 2021), the prevalence of mixed infections was significantly higher in females compared to males and this could be due to the maternal transmission (Ahmad et al., 2017), ineffective transmission of one strain (Kittayapong et al., 2002), or physiological mechanisms allowing *Wolbachia* infection in females but not in males (Kittayapong et al., 2002). The exact mechanisms of *Wolbachia* infection and dissemination in mosquito vectors remain unclear (Bi & Wang, 2020).

The prevalence of *w*AlbB infection was higher than *w*AlbA, supporting prior findings for European populations (Tortosa et al., 2010; Shaikevich et al., 2019; Belo, 2021). This inequality is particularly pronounced in males and could be attributed to reduced *w*AlbA density over time (Shaikevich et al., 2019) or a lower reproduction rate (Kittayapong et al., 2002; Wiwatanaratanabutr & Kittayapong, 2009). Research has suggested that females may not efficiently transmit *w*AlbA to their progeny, resulting in a reduction in male fertility due to Cytoplasmic Incompatibility (CI) (Tortosa et al., 2010).

In this study, males from São Tomé population were not infected by *Wolbachia*. In fact, low *Wolbachia* prevalence has been detected in natural *Ae. albopictus* infections in Cameroon, one of the possible origins of *Ae. albopictus* from São Tomé (Bamou et al., 2021). Therefore, is not possible to assume that males in this region do not host *Wolbachia*, as infected females were found. Thus, the absence of infected males in São Tomé may be related to the lower prevalence of endosymbiont infection in males and the small sample size (N=2).

The number of *Wolbachia*-negative samples was minimal (1.11%), indicating that innate *Wolbachia* infection may provide selective advantages to hosts, such as higher fecundity or hatching rates (Dobson et al., 2004). Geographic variation in *Wolbachia* infection has been reported (Joanne et al., 2015), but this study did not find significant differences between samples from different locations in Spain, as well as other studies carried out in other locations of this country (Bueno-Marí et al., 2023). A uniform distribution pattern with a predominance of mixed infections has been suggested and supported by prior studies for *Ae. albopictus* populations across Europe (Belo, 2021).

The rapid dissemination and uniformity of distribution in Europe could result from multiple introductions (Kotsakiozi et al., 2017; Pichler et al., 2019) in different Mediterranean countries, leading to constant mixed infection of *Ae. albopictus* by *Wolbachia.* Maternal inheritance and cytoplasmic incompatibility may also play a role in promoting the spread of *Wolbachia* infection (Alphey, 2014).

### Aedes albopictus vector control

To control *Ae. albopictus* populations, one approach involves creating a colony with a stable infection of specific exogenous *Wolbachia* strains that cause Cytoplasmic Incompatibility (CI) and non-viable offspring. The Incompatible Insect Technique (IIT) involves the release of males carrying these *Wolbachia* strains into the environment, leading to a reduction in the target population. This approach has been tested and implemented in *Ae. albopictus* populations which naturally hosted two *Wolbachia* strains (*w*AlbA and *w*AlbB). Introducing a third strain, *w*Pip (found in *Culex pipiens molestus*), resulted in a triple-infected colony that effectively decreased the number of *Ae. albopictus* females in the study area (Mains et al., 2016). Recent strategies employed *w*Pip transinfection in *Ae. albopictus*, accompanied by the removal of its native *Wolbachia* strains, yielding promising outcomes (Caputo et al., 2020). These findings underscore IIT’s potential as a future tool for *Ae. albopictus* control.

To conclude, this study significantly enhances our understanding of the bacteriome of *Ae. albopictus* populations from Spain and São Tomé. By unveiling the presence of *Wolbachia* and other bacterial endosymbionts, we establish a foundational framework for future manipulations of endosymbionts and the introduction of new strains for vector control. As the scientific community intensifies its search for innovative and effective methods to combat vector-borne diseases, the results of this study become increasingly valuable.

Future research work to expand upon our findings include: (i) examine the impact of the bacteriome on *Ae. albopictus* vector competence to better understand how specific bacterial endosymbionts affect the mosquito’s ability to survive and transmit arboviruses. This could lead to the development of predictive models for better disease management; (ii) based on our knowledge of the microbiome, evaluate the potential of CRISPR/Cas9 gene editing in endosymbionts for vector control, focusing on genes that affect vector competence and reduce pathogen transmission; (iii) investigate paratransgenic strategies by introducing recombinant bacteria into the mosquito microbiome to decrease pathogen transmission and address insecticide resistance; (iv) assess the ecological consequences of microbiome manipulation, considering long-term effects on mosquito populations, non-target organisms, and ecosystems to ensure sustainability; (v) use advanced sequencing technologies such as nanopore sequencing will provide more comprehensive and precise 16S gene sequences enabling a more detailed understanding of the mosquito microbiome and its implications for vector control.

By pursuing these research lines, we can advance medical and veterinary entomology and make significant progress in controlling vector-borne diseases, building upon the findings of this study to have a significant global health impact.

### Aedes albopictus vector control

To control *Ae. albopictus* populations, one approach involves creating a colony with a stable infection of specific exogenous *Wolbachia* strains that cause Cytoplasmic Incompatibility (CI) and non-viable offspring. The Incompatible Insect Technique (IIT) involves the release of males carrying these *Wolbachia* strains into the environment, leading to a reduction in the target population. This approach has been tested and implemented in *Ae. albopictus*, which naturally hosts two *Wolbachia* strains (*w*AlbA and *w*AlbB). Introducing a third strain, *w*Pip (found in *Culex pipiens molestus*), resulted in a triple-infected colony that effectively decreased the number of *Ae. albopictus* females in the study area (Mains et al., 2016). Recent strategies employed *w*Pip transinfection in *Ae. albopictus*, accompanied by the removal of its native *Wolbachia* strains, yielding promising outcomes (Caputo et al., 2020). These findings underscore IIT’s potential as a future tool for *Ae. albopictus* control.

To conclude, this study significantly enhances our understanding of the bacterial microbiome of *Ae. albopictus* populations from Spain and São Tomé. By unveiling the presence of *Wolbachia* and other bacterial endosymbionts, we establish a foundational framework for future manipulations of endosymbionts and the introduction of new strains for vector control. As the scientific community intensifies its search for innovative and effective methods to combat vector-borne diseases, the insights from this study become increasingly invaluable.

Future research directions to expand upon our findings include: (i) examine the impact of the bacterial microbiome on *Ae. albopictus* vector competence to better understand how specific bacterial endosymbionts affect the mosquito’s ability to survive and transmit arboviruses. This could lead to the development of predictive models for better disease management; (ii) based on our knowledge of the microbiome, evaluate the potential of CRISPR/Cas9 gene editing in endosymbionts for vector control, focusing on genes that affect vector competence and reduce pathogen transmission.; (iii) investigate paratransgenic strategies by introducing recombinant bacteria into the mosquito microbiome to decrease pathogen transmission and address insecticide resistance; (iv) assess the ecological consequences of microbiome manipulation, considering long-term effects on mosquito populations, non-target organisms, and ecosystems to ensure sustainability; (v) use advanced sequencing technologies such as nanopore sequencing will provide more comprehensive and precise 16S gene sequences enabling a more detailed understanding of the mosquito microbiome and its implications for vector control.

By pursuing these research directions, we can advance medical and veterinary entomology and make significant progress in controlling vector-borne diseases, building upon the findings of this study to have a significant global health impact.

## Acknowledgments

This work was initially conceived and designed during research stay of DBB at GHTM in 2019, as part of the Short-Term Scientific Missions (STSM) within the AIM-COST CA17108 Action. Subsequently, this study was funded by the Fundação para a Ciência e Tecnologia funding to GHTM-UID/04413/2020, LA-REAL-LA/P/0117/2020 and “healTh RIsk and social vulnerability to Arboviral Diseases in mainland Portugal” (TRIAD) – Ref. PTDC/GES-OUT/30210/2017.

