## Supplementary material for "Characterization of the microbiome of *Aedes albopictus* populations in different habitats from Spain and São Tomé: implications for vector control": Supp. Table S1

### Supplementary tables

**Table S1 -** Description of primers used in *Wolbachia* genotyping.

| | Primer Forward (Fw) | Primer Reverse (Rv) | Size of product |
|---|---|---|---|
| ***w*AlbA** | 328F:5'-CCAGCAGATACTATTGCG-3' | 691R:5'-AAAAATTAAACGCTACTCCA - 3' | 389 bp |
| ***w*AlbB** | 183F:5’-AAGGAACCGAAGTTCATG - 3’ | 691R:5'-AAAAATTAAACGCTACTCCA-3' | 501 bp |
| ***w*sp** | 81F:5’-TGGTCCAATAAGTGATGAAGAAAC –3’ | 691R:5'-AAAAATTAAACGCTACTCCA-3' | 615 bp |

**Table S2 -** Correlation between the number of OTUs and the diversity of the data set, in the samples from Sao Tome. 1) Input sequences. 2) Sequences after preprocessing and chimera removal. 3) Sequences assigned to OTUs. 4) Sequences assigned to taxa. 5) Count after lineage-specific copy-number correction. 6) Median sequence length after pre-processing.

| Sample | 1 | 2 | 3 | 4 | 5 | 6 |
|---|---|---|---|---|---|---|
| **STPF0.1** | 125 964 | 100.0% | 88.7% | 88.7% | 102 138 | 399 |
| **STPF0.2** | 105 677 | 100.0% | 82.9% | 82.9% | 74 788 | 399 |
| **STPF0.3** | 155 388 | 100.0% | 88.4% | 88.4% | 126 113 | 399 |
| **STPF0.4** | 122 065 | 100.0% | 87.2% | 87.2% | 97 713 | 399 |
| **STPF0.5** | 97 783 | 100.0% | 84.5% | 84.5% | 74 320 | 399 |

**Table S3-** Correlation between the number of OTUs and the diversity of the data set, in the samples from Spain. 1) Input sequences. 2) Sequences after pre-processing and chimera removal. 3) Sequences assigned to OTUs. 4) Sequences assigned to taxa. 5) Count after lineage-specific copy-number correction. 6) Median sequence length after pre-processing.

| Sample | 1 | 2 | 3 | 4 | 5 | 6 |
|---|---|---|---|---|---|---|
| **MZF0.1** | 53 154 | 100.0% | 82.4% | 82.4% | 40 146 | 399 |
| **MZF0.2** | 56 715 | 100.0% | 82.1% | 82.1% | 42 741 | 399 |
| **MZF0.3** | 58 170 | 100.0% | 78.9% | 78.9% | 41 120 | 399 |
| **MZF0.4** | 59 498 | 100.0% | 82.5% | 82.5% | 45 081 | 399 |
| **MZF0.5** | 56 249 | 100.0% | 79.8% | 79.8% | 40 865 | 399 |
| **PAF0.1** | 62 949 | 100.0% | 80.3% | 80.3% | 46 164 | 399 |
| **PAF0.2** | 65 701 | 100.0% | 81.2% | 81.2% | 49 022 | 399 |
| **PAF0.3** | 56 048 | 100.0% | 80.6% | 80.6% | 41 421 | 399 |
| **PAF0.4** | 43 783 | 100.0% | 78.4% | 78.4% | 31 255 | 399 |
| **RCF0.1** | 65 827 | 100.0% | 81.5% | 81.5% | 49 296 | 399 |
| **RCF0.2** | 43 868 | 100.0% | 77.2% | 77.2% | 31 177 | 399 |
| **RCF0.3** | 65 500 | 100.0% | 81.4% | 81.4% | 48 967 | 399 |
| **RCF0.4** | 58 493 | 100.0% | 82.4% | 82.4% | 44 330 | 399 |
| **RCF0.5** | 52 145 | 100.0% | 79.2% | 79.2% | 37 996 | 399 |

**Table S4 -** Average, per sample, of the Shannon and Simpson indices for the Spain and São Tomé populations.

| Sample | Shannon (average) | Simpson (average) |
|---|---|---|
| **MZF0.1** | 4,388 | 0,822 |
| **MZF0.2** | 4,327 | 0,816 |
| **MZF0.3** | 4,788 | 0,851 |
| **MZF0.4** | 4,342 | 0,814 |
| **MZF0.5** | 4,289 | 0,799 |
| **PAF0.1** | 4,579 | 0,820 |
| **PAF0.2** | 4,545 | 0,837 |
| **PAF0.3** | 4,307 | 0,815 |
| **PAF0.4** | 4,246 | 0,814 |
| **RCF0.1** | 4,364 | 0,821 |
| **RCF0.2** | 4,353 | 0,820 |
| **RCF0.3** | 3,999 | 0,804 |
| **RCF0.4** | 4,654 | 0,836 |
| **RCF0.5** | 4,243 | 0,791 |
| **STPF0.1** | 4,010 | 0,771 |
| **STPF0.2** | 4,403 | 0,826 |
| **STPF0.3** | 3,411 | 0,659 |
| **STPF0.4** | 4,131 | 0,790 |
| **STPF0.5** | 4,400 | 0,818 |
